# Application of cryo-FIB-SEM for investigating organelle ultrastructure in guard cells of higher plants

**DOI:** 10.1101/2024.08.30.610476

**Authors:** Bastian Leander Franzisky, Xudong Zhang, Claus Jakob Burkhardt, Endre Majorovits, Eric Hummel, Andreas Schertel, Christoph-Martin Geilfus, Christian Zörb

## Abstract

Stomata are vital for CO2 and water vapor exchange, with guard cells’ aperture and ultrastructure highly responsive to environmental cues. However, traditional methods for studying guard cell ultrastructure, which rely on chemical fixation and embedding, often distort cell morphology and compromise membrane integrity, leaving no suitable methodology until now. In contrast, plunge-freezing in liquid ethane rapidly preserves cells in a near-native vitreous state for cryogenic electron microscopy. Using this approach, we applied Cryo-Focused Ion Beam-Scanning Electron Microscopy (cryo-FIB-SEM) to study the guard cell ultrastructure of *Vicia faba*, a higher plant model chosen for its sensitivity to external factors and ease of epidermis isolation, advancing beyond previous cryo-FIB-SEM applications in lower plant algae. The results firstly introduced cryo-FIB-SEM volume imaging, enabling subcellular ultrastructure visualization of higher plants like *V. faba* in a vitrified, unaltered state. 3D models of organelles such as stromules, chloroplast protrusions, chloroplasts, starch granules, mitochondria, and vacuoles were reconstructed from cryo-FIB-SEM volumetric data, with their surface area and volume initially determined using manual segmentation. Future studies using this near-native volume imaging technique hold promise for investigating how environmental factors like drought or salinity influence stomatal behavior and the morphology of guard cells and their organelles.

## 1. Introduction

Stomata, composed of paired guard cells either in kidney shapes or dumbbell shapes in all vascular plant species,[1] serve as fundamental gatekeepers regulating the exchange of atmospheric CO2 and H2O, particularly vital in challenging conditions such as drought and soil salinity.[2] Environmental factors such as these can alter stomatal aperture, density, number, size, and morphology.[3] For example, when the stomatal opening drives the influx of CO2 for photosynthesis, stomatal closure is essential to diminish water loss when plants are exposed to drought stress. In maize crops, reduced soil water content significantly increased stomatal amount, leading to an increase in stomatal density but a decrease in stomatal size and aperture. A negative correlation was found between stomatal density and photosynthetic rate.[4, 5] By contrast, a positive correlation between transpiration rate and stomatal density have been reported in barley and wheat under soil salinity, where stomata size was also affected.[6] Obviously, external cues significantly influence stomatal behavior, but how the ultrastructures of guard cells are affected and respond to the environmental stimuli remains unknown. Investigating the ultrastructural variations of organelles at the subcellular levels are of vital importance to understand acclimatory mechanisms of plants to adverse environments. The presented methodological work firstly applied the cryo-Focused Ion Beam (FIB)-Scanning Electron Microscope (SEM) technique on guard cells to visualize subcellular organelles and conduct a tentative quantification of organelle size. Up to now, quantifying stomata morphology traits have been widely applied through analyzing the collected microscopic images from light microscopes or scanning electron microscopes to calculate stomatal density and size either in a manual way or in an algorithm-based machine learning process.[7, 8] In this context, the samples are usually intact fresh leaves attached to the plant[3, 5] or cut off,[6] however, the samples are not readily accessible for ultrastructural analysis. The crucial first step is the sample preparation, which directly affects the integrity of the sample and thus the reliability of the image data obtained. A particular challenge is to prepare plant samples for subsequent microtomic dissection of individual cells without affecting the native state of the cells, which can be a significant hurdle and might be responsible for the lack of studies of guard cell ultrastructure.

Conventional electron microscopy (EM) sample preparation involves chemical treatments like glutaraldehyde or osmium tetroxide to fix plant tissues.[9, 10] Given the high water content inherent in plant cells, significant challenges arise, including fixation artifacts such as membrane ruptures during chemical fixation.[11, 12] Moreover, diffusion of fixatives and resins is impeded by intercellular air spaces, cell walls, and hydrophobic surfaces like waxes (e.g., cuticles of stomata), exacerbating limitations of chemical fixation such as time-consuming diffusion and selectivity of fixatives.[13] Also inherent staining with heavy metals for achieving imaging contrast in resin-embedded samples poses its own challenges, including dehydration and deformation of plant specimens and resolution limited by staining grain size.[14] Consequently, preserving plant ultrastructural features close to their natural state, using resin-embedding preparation protocols for electron microscopy, remains exceedingly challenging. Any chemical treatment, prior to or during chemical fixation, are stressors triggering a response resulting in altered subcellular ultrastructure and eliminates any investigation how environmental factors influence plant behavior. For example, in living plant cells, intracellular water may physiologically be lost due to high salt concentration in phosphate-buffered saline (PBS) solution, potentially influencing internal structure and membrane integrity.

In contrast, cryo-immobilization by plunge or high pressure freezing allows subcellular ultrastructure visualization in the near native vitrified state by electron microscopy at cryogenic conditions.[15, 16] While sample vitrification followed by freeze substitution and resin embedding permits room temperature electron microscopy (EM) investigation, it is still susceptible to resin embedding related artifacts like mentioned above. Plunge-freezing provides reliable vitrification of biological samples to a depth of up to 10 µm,[17] whereas high-pressure freezing yields good vitrification up to a specimen thickness of 5 to 200 µm.[18]

The FIB technology enables advanced single-cell cross-sectioning and volume imaging of subcellular organelles. The FIB-milling lamella preparation method for cryo-electron tomography (cryoET) imaging and subtomogram averaging has been extensively developed in both eukaryotic and bacterial model organisms.[19–21] This technique has successfully revealed the 3D structural architecture of chloroplast cells using cell cultures of the aquatic alga Chlamydomonas.[22, 23] Besides, cryo-TEM imaging of knife-cut ultrathin sections (CEMOVIS) can provide high-resolution 3D data,[24] but the restricted field of view is limiting the accessible volume. By contrast, serial SEM images obtained from FIB-milling slices in the FIB-SEM system can generate large volume data, although the image resolution is diminished in comparison with TEM. Relying on the sample preparation method of high pressure freezing and freeze substitution, FIB-SEM has been applied on wild Chlorella pyrenoidosa cells, clearly distinguishing between mitochondria, plastids, Golgi, and other membranes such as the endoplasmic reticulum (ER) and tonoplast based on resin-embedded samples at room temperature[25] and mouse optic nerves cells and Bacillus subtilis spores visualizing Golgi cisternae, vesicles, ER, and mitochondrial cristae under cryogenic state.[17] More importantly, the FIB-SEM technique allows 3D visualization after reconstructing the serial image stack, which represents consecutive images of sample sections.

Leaf stomata have often been studied using scanning electron microscopy (SEM), either with alive leaf tissues or with common methods including chemical fixations such as glutaraldehyde, phosphate-buffered saline (PBS) solution, and ethanol.[3, 5–8] Typically, whole plant leaves are mounted for analysis rather than isolated leaf epidermis, primarily due to the challenges associated with obtaining pure epidermis for microscopy. The leaf epidermis presents a complex specimen for fixation, comprising various cell types including primary epidermal cells, trichomes, papillae, and guard cells forming the stomata and subsidiary cells. Additionally, the epidermal tissue is typically covered by outer layers of cuticle and/or waxes, further complicating fixation procedures.[26] This has led to a lack of detailed knowledge regarding guard cell ultrastructure, as studying the whole plant leaf tissue at the subcellular level proves challenging. To address this gap, we utilized *V. faba* as a model crop for higher plants to explore the potential of cryo-FIB-SEM volume imaging. This technique allows for the visualization of 3D ultrastructure and identification of subcellular organelles for surface area and volume quantification based on peeled epidermis in a near-to-native state.

This study firstly introduces the use of cryo-FIB-SEM techniques to analyze leaf cells of vascular plants, specifically focusing on guard cells and stomata (Figure 1). We isolated a monolayer of plant cells, enriched with guard cells and stomata from *V. faba*, for cryogenic plunge freezing and FIB-SEM analysis. This approach represents a significant advancement over studies on liquid-cultured chloroplast cells from lower plant algae.[22, 23] Compared to resin preparation, cryo-preparation in electron microscopy more effectively preserves native structures, lipid membranes, and chemical composition, ensuring high-resolution imaging with enhanced safety and efficiency.[27] In future studies, this advanced method offers a promising new approach for exploring plants’ responses to stressors such as drought and salt, providing valuable insights into the environmental influences on guard cell ultrastructure and organelle size.

**Figure 1.**
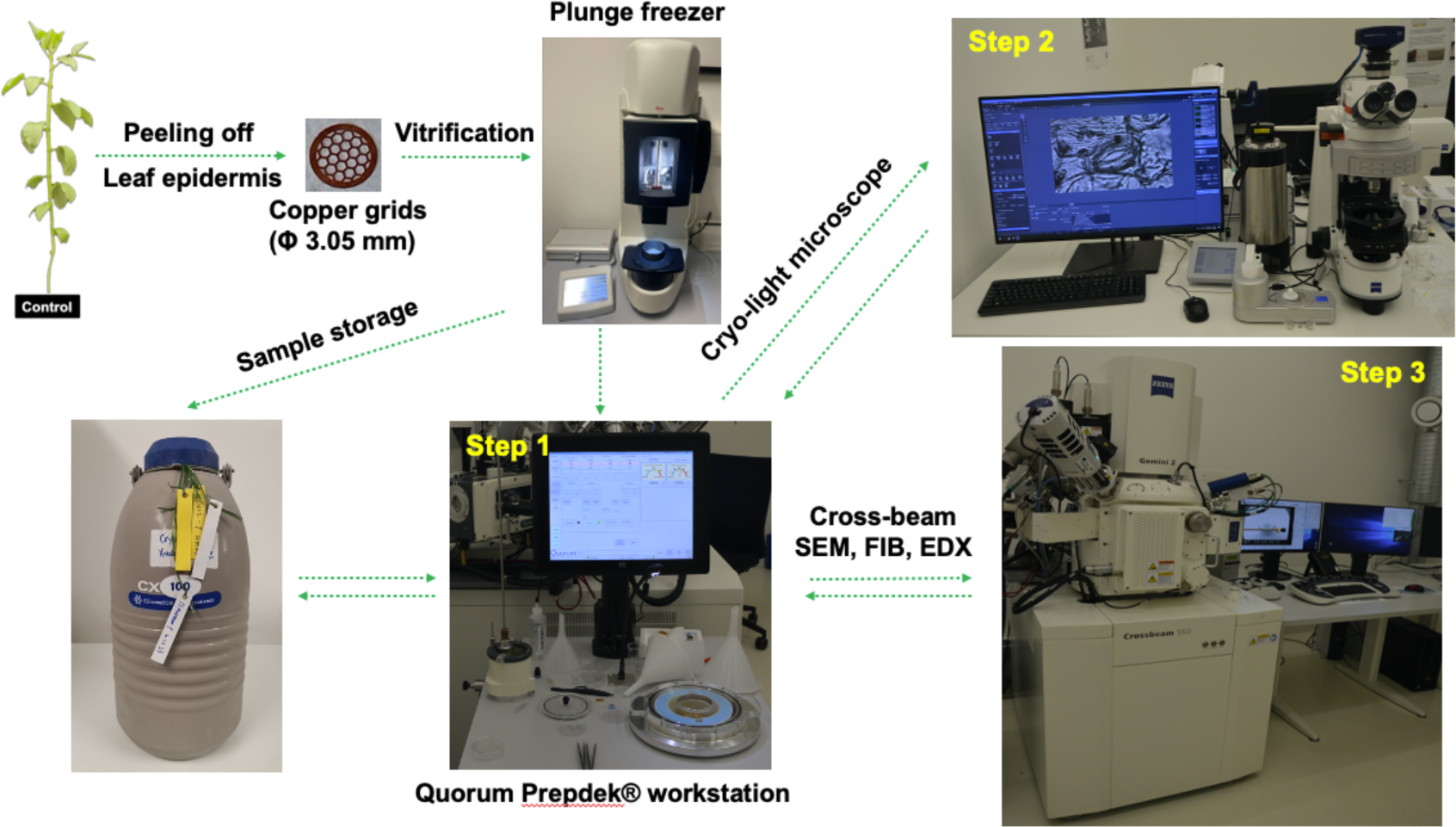
**The cryo-LM and FIB-SEM workflow for observing the ultrastructure of guard cells in *V. faba***. Preparation of the plant sample for plunge freezing with medium blotting force for 2-5 seconds in liquid ethane (Leica EM GP2). Transfer of the vitrified sample to the Quorum loading station (Step 1) where the sample is loaded into the correlative cryo holder. The loading station minimizes the risk of sample damage or contamination, because the TEM grid needs to be mounted only once for all imaging experiments, minimizing the risk of additional ice contamination and potential sample damage from handling with forceps during a second mounting process. Transfer into the Zeiss LSM900 equipped with a Linkam cryo-stage for cryo-LM (Step 2) followed by correlative FIB-SEM data acquisition in the Zeiss cryo-FIB-SEM Crossbeam 550 (Step 3). Intermediate sample storage in a liquid nitrogen dewar is possible at all times.

## 2. Materials and Methods

### 2.1. Plant cultivation

The *Vicia faba* L. variety Fuego (obtained from Norddeutsche Pflanzenzucht Hans-Georg Lembke KG, Hohenlieth, Germany) was cultivated under hydroponic conditions within a climate cabinet (WEISS HGC1014, Heuchelheim, Germany), with a 14/10- hour day/night cycle, temperatures maintained at 22/18°C, and humidity levels around 80/60%. Light intensity was approximately 300 μmol photons m^2^/s at shoot level.[28] Initially, seeds were soaked in aerated CaSO4 solution (0.5 mM) for 1 day at room temperature, then transferred to moistened quartz sand. After 12 days of germination, seedlings were transplanted into plastic pots filled with 1/4-strength aerated nutrient solution. The nutrient concentration was gradually increased: to 1/2-strength after 2 days, 3/4-strength after 3 days, and full-strength after 4 days. The composition of the full-strength nutrient solution included: 0.1 mM KH2PO4, 1.0 mM K2SO4, 2.0 mM Ca(NO3)2, 0.5 mM MgSO4, 0.00464% (wt/vol) Sequestren (Ciba Geigy, Basel, Switzerland), 10 μM NaCl, 10 μM H3BO3, 2.0 μM MnSO4, 0.5 μM ZnSO4, 0.2 μM CuSO4, 0.1 μM CoCl2, and 0.05 μM (NH4)6Mo7O24. Plants were harvested 27 days after reaching full nutrition strength.

### 2.2 Specimen preparation and plunge freezing

At the harvest stage, the abaxial leaf epidermis with a rough thickness of 7 µm was delicately peeled off using forceps. This sample thickness is compliant with a vitrification depth of around 10 µm when plunge freezing the specimen.[17] The epidermal fragments of 4 to 6 mm^2^ were then floated on tap water at room temperature and promptly mounted onto hexagonal support-free 50 mesh TEM copper grids purchased from ScienceServices. (see Figure 1). Excess water was gently removed from the grids by blotting them carefully with filter paper. Subsequently, the grids were placed into a plunge freezer (EM GP2, Leica Microsystems, see Figure 1). After another round of semi-automated blotting with filter paper in the EM GP2 plunge freezer (standardized procedure with 2-5 seconds blotting time), the grids were plunge-frozen in liquid ethane. In total, five sample grids were plunge-frozen and then transferred into a grid box and stored in liquid nitrogen until microscopic analysis.

### 2.3 Correlative cryo LM and FIB-SEM

The acquisition of correlative cryo LM and FIB-SEM data was performed using a cryo-system consisting of the following three hardware components: the loading station (Quorum Technologies, modified by Carl Zeiss Microscopy GmbH) for loading and transferring cryo samples, see Figure 1 Step 1, the LSM 900 light microscope (Carl Zeiss Microscopy GmbH, Oberkochen, Germany) equipped with a Linkam cryo stage (Linkam Scientific, Salfords, UK), see Figure 1 Step 2, and the FIB-SEM Crossbeam 550 (Carl Zeiss Microscopy GmbH) equipped with a Quorum cryo stage (Quorum Technologies, Laughton, UK), see Figure 1 step 3.

The loading station facilitates easy insertion of the TEM grids into the correlative cryo holder. Once inserted, the sample remains mounted on the correlative cryo holder and can be seamlessly transferred onto corresponding adaptors for LM or FIB-SEM observation (as depicted by details of loading station in Figure S1). This setup ensures that the grids are mounted only once for all measurements, which helps to maintain consistent sample orientation, and eliminates the risk of additional ice contamination and of damaging the sample with the forceps due to a second mounting process. After LM or SEM imaging, the correlative cryo holder can be stored under liquid nitrogen until the next imaging session (Figure 1).

The ZEN acquisition and visualization software (Carl Zeiss Microscopy GmbH) was employed for correlative LM and SEM data acquisition. Following the LM to FIB-SEM three-point-alignment to register the two stage coordinate systems, the LM dataset facilitates navigation to the areas of interest and acquisition of 3D FIB-SEM datasets. The ZEN image processing toolbox allows to align the image stacks and to denoise the images as described in detail in chapter 2.6. The software Dragonfly (Object Research Systems Inc, Canada) was used for the segmentation of cell organelles, 3D visualization and quantification.

### 2.4 LM data acquisition

LM images were acquired with an LSM 900 light microscope (Carl Zeiss Microscopy GmbH, Oberkochen, Germany) equipped with a Linkam cryo stage (Linkam Scientific, Salfords, UK). After cooling down the Linkam cryo stage to liquid nitrogen temperature, the correlative cryo holder was transferred onto the stage and the sample was imaged in bright field mode at 5x magnification to check the sample integrity and to exclude that the sample is covered with a thick ice layer (>> 1 µm), see Figure 2 A. Subsequently, the sample was imaged at different magnifications (20x and 100x) in bright field mode (Figure 2 B and C) and in fluorescence mode at wavelength ranges of 615 nm to 648 nm for chlorophyll (Figure 2 D) and 450 nm to 488 nm for autofluorescence from cell walls (Figure 2 E).

**Figure 2.**
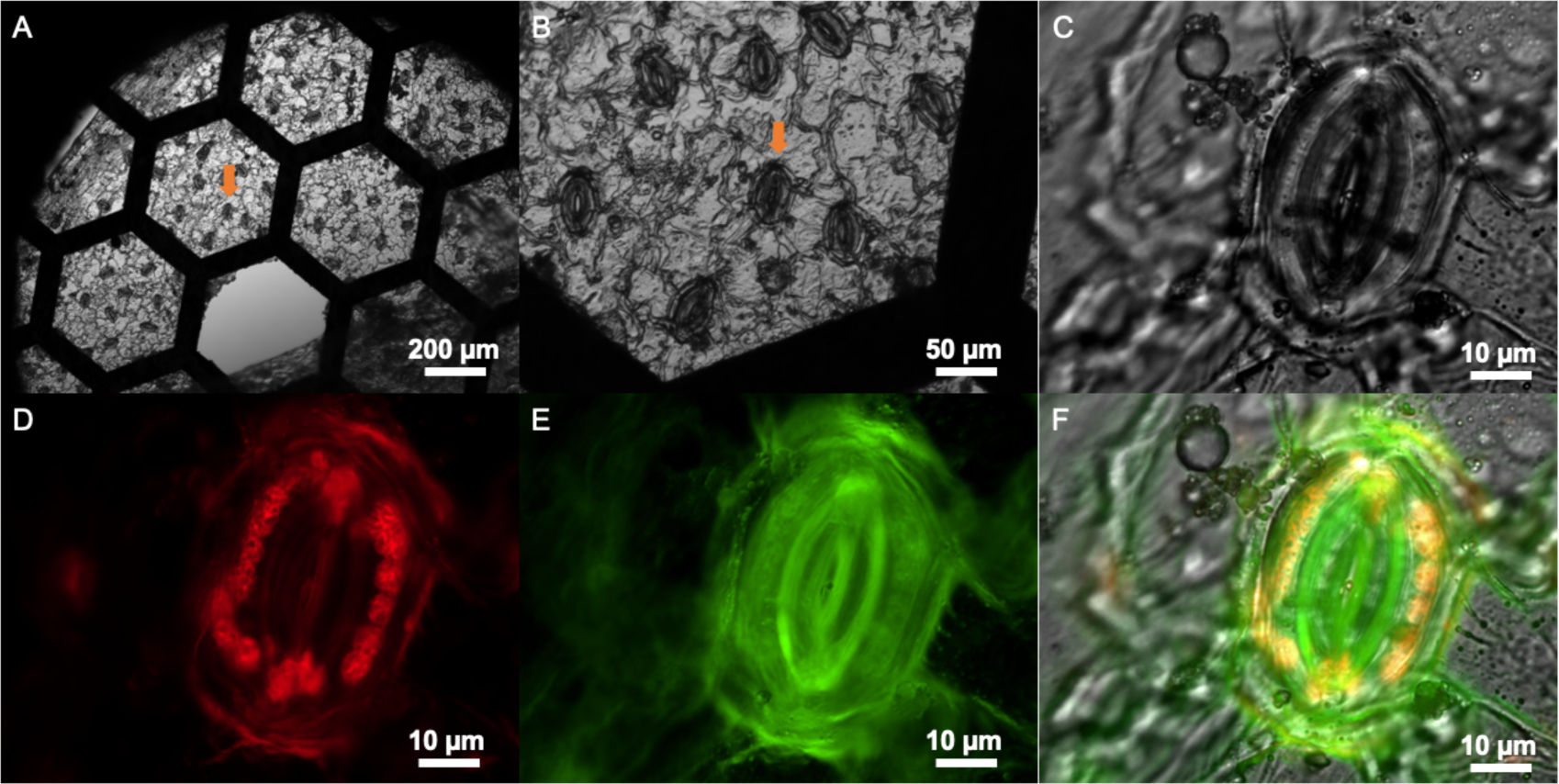
Bright-field and fluorescence imaging of guard cells using cryo-LM. A-C : Bright field images with different magnifications (A: 5x, B: 20x, C: 100x), orange arrows indicated the target guard cell. D and E: Fluorescence LM images acquired at a magnification of 100x. D: Chlorophyll autofluorescence signal (maximum at 638 nm, ranging from 615 nm to 648 nm) showing chloroplasts (displayed in red). E: Cell wall signal (maximum at 475 nm, ranging from 450 nm to 488 nm) showing in green. F: Merged image of the three LM channels C-E.

### 2.5 Sputter coating and FIB-SEM data acquisition

Inside the Quorum prep chamber which is attached to the FIB-SEM and serves as an airlock and a sample preparation tool at the same time (options for sputter coating, sublimation and freeze fracture), the sample was sputter coated to make the surface conductive thereby minimizing charging artifacts during SEM imaging (Figure 3 B and D). Sputter coating was performed using platinum with a 5 mA current for 45 seconds to produce a platinum layer of about 40 nm thickness.

**Figure 3.**
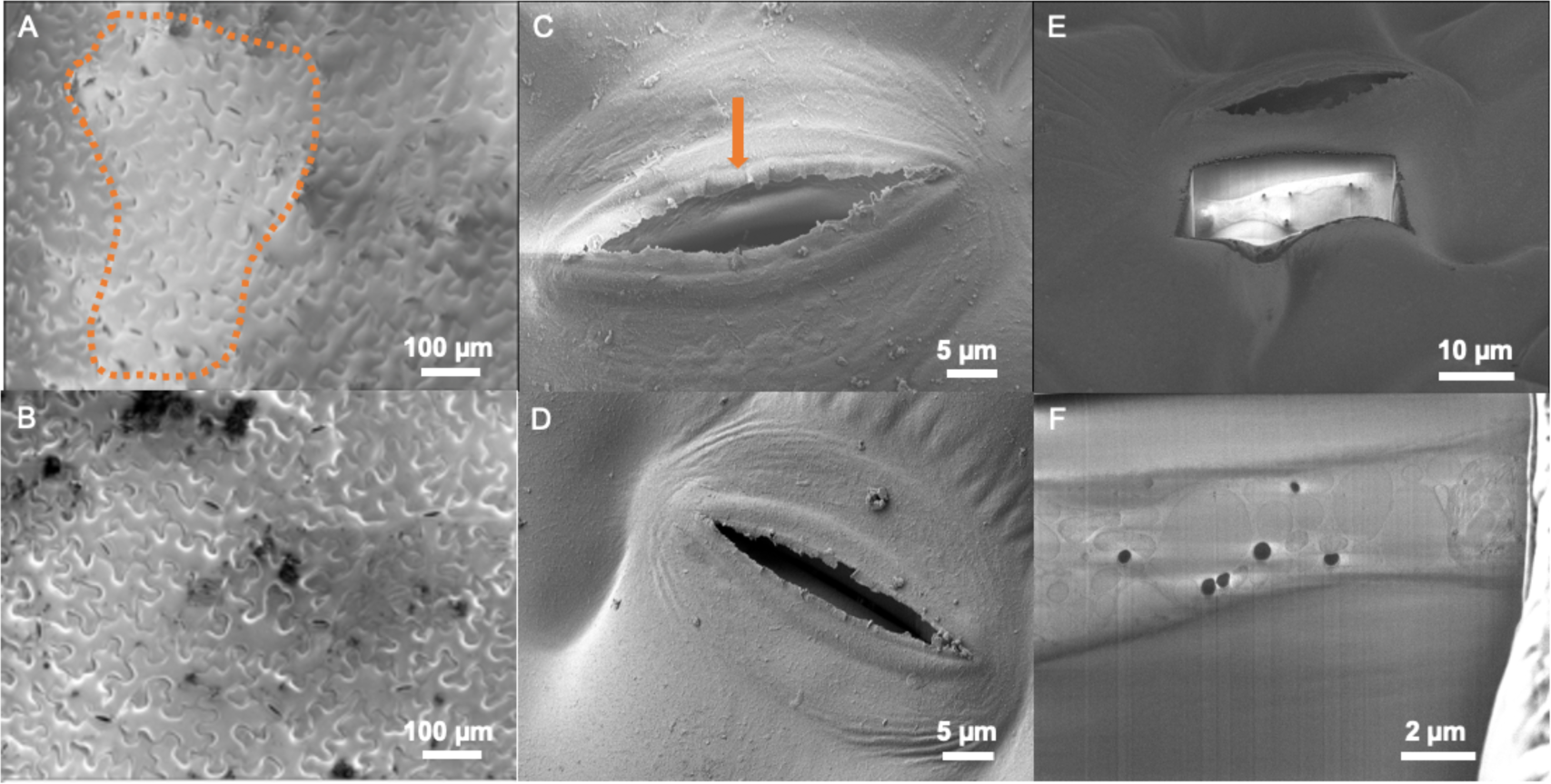
SEM imaging of the sample surface and 3D FIB-SEM data collection. A-D : Imaging before and after platinum sputter coating to reduce charging effects. A: Charging artifacts encircled by an orange dotted line are visible over the waxy surface of the epidermis. B: Charging artifacts are reduced after sputter coating. C: stomata imaged without sputter coating, charging clearly visible on the rims of the stomata as indicated by orange arrow. D: Charging artifacts minimized after a short sputter coating cycle. E: 25 µm wide trench milled with a 7 nA FIB current for FIB-SEM imaging to investigate ultrastructure of guard cell organelles. F: Raw image slice from the 3D FIB- SEM dataset (SEM InLens SE detector, 2.0 kV, 30 pA) to visualize different organelle regions at the cross section interface after FIB milling.

After transferring the sputter coated sample onto the Quorum cryo stage, the LM to FIB-SEM stage registration was performed within the ZEN user interface based on the previously acquired LM dataset. Subsequently, areas of interest were relocated and cross-sections were prepared for serial cryo-FIB sectioning and SEM imaging. Initial trenches were milled with a FIB current of 7 nA at 30 kV, cross-section polishing and volume imaging were performed with a 700 pA FIB current. The slice thickness was set to 20 nm. The SEM images used for image processing and data analysis were acquired with a pixel size of 11.3 nm, an EHT of 2.3 kV and a SEM beam current of 30 pA using the InLens SE detector. The resulting 3D FIB-SEM dataset had a volume of 23.1 µm x 17.4 µm x 6.4 µm (image size: 23.1 µm x 17.4 µm; milling depth: 6.4 µm).

### 2.6 FIB-SEM data processing

After acquiring the 3D FIB-SEM datasets, the images underwent alignment and processing using various post processing options provided by the ZEN "EM processing toolbox". The SEM images acquired during the volume imaging experiments are affected by the so-called curtaining effect (Figure 4A), vertical stripes that result from uneven milling due to particles on the sample surface and density variations within the sample volume. These stripes were effectively removed using the “Remove stripes” method in the ZEN "EM processing toolbox" which is based on the VSNR (Variational Stationary Noise Remover) algorithm.[29] This algorithm removes stripes from images in arbitrary directions and acts as a denoising filter at the same time, resulting in improved image quality (Figure 4B). ZEN software’s destriping function combines smoothing and contrast enhancement to reduce striping artifacts and improve image quality. Smoothing, which includes spatial and frequency domain filtering, reduces noise and clarifies main structures. Contrast enhancement improves feature visibility using techniques such as adaptive histogram equalization and intensity transformations. The process involves transforming the image to the frequency domain using Fourier Transform, where periodic noise appears as distinct peaks that are then filtered out. The filtered image is converted back to the spatial domain, and adaptive filtering applies smoothing and contrast enhancement based on local image characteristics to preserve structures while reducing noise.

**Figure 4.**
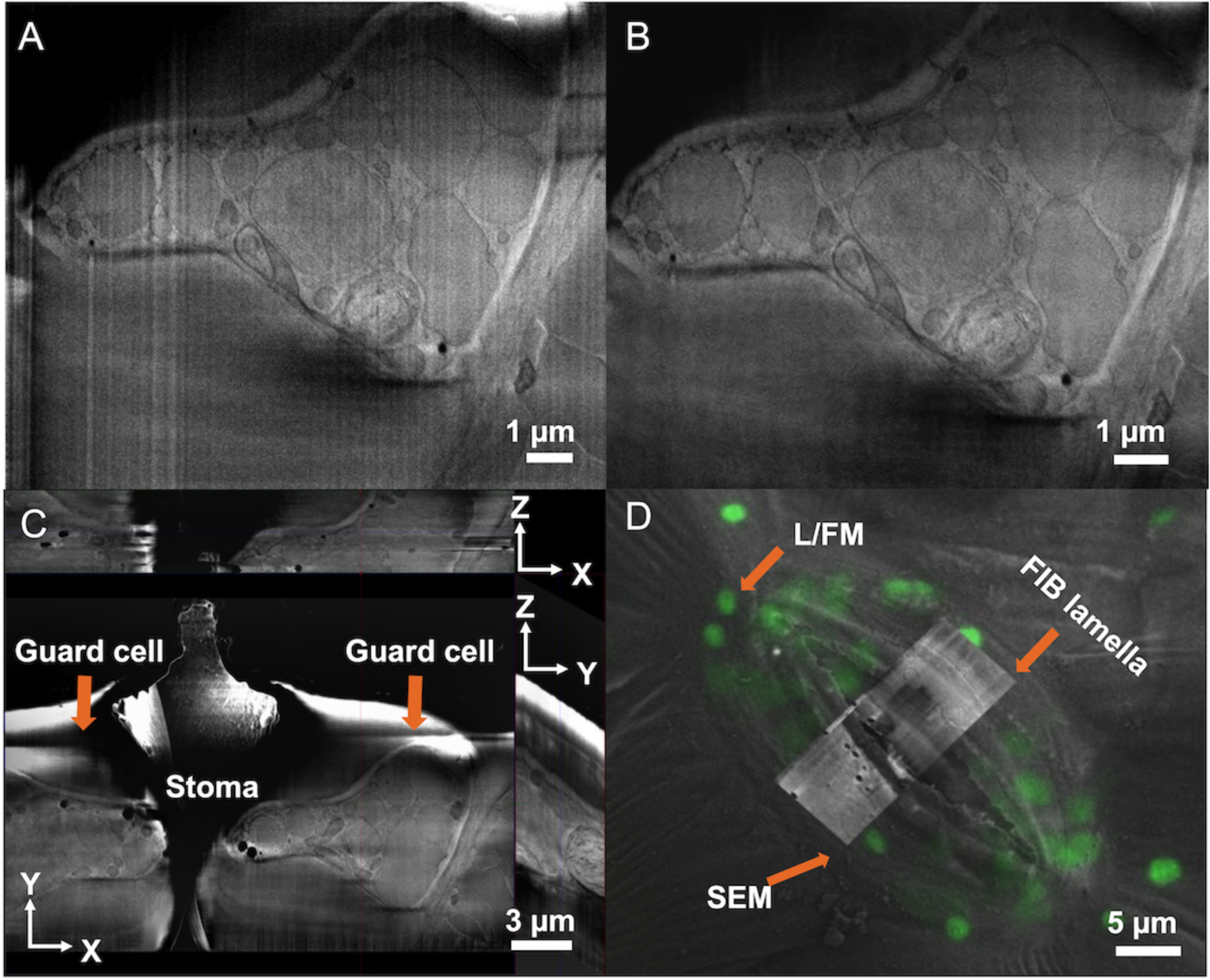
Post-processing and visualization of 3D FIB-SEM dataset. A: Raw image from 3D FIB-SEM dataset showing typical milling artifacts (curtaining effect). B: Processed image: the destriping tool (VSNR algorithm) reduces the milling artifacts and enhances feature visibility. C: Orthoview showing the processed 3D FIB- SEM dataset as perpendicular planes (XY, XZ and YZ). D: Overlay of light/fluorescent images (L/FM), SEM image and FIB lamella.

Curtaining poses challenges when measuring specimen volumes. Our study did not directly compare different curtaining levels. However, literature suggests destriping does not significantly affect quantification.[30–33] In comparison with ZEN’s destriping function, which uses Fourier Transform and adaptive filtering, IMOD Software[33] employs similar principles, utilizing Fourier Transform and median filtering to effectively manage stripe artifacts, confirming that these artifacts can be handled without impacting quantification. While our approach is supported by literature, empirical validation is needed in future studies. Additionally, the ZEN software allowed for result visualization using Ortho- and 3D view options (Figure 4 C). A stack of 320 images generated by FIB-SEM volume imaging was processed using the visualization and processing software Dragonfly (2020.2; Object Research Systems Inc) to create a 3D model of spatial organelle arrangement.

Manual segmentation was employed in this study due to the complexity of identifying organelles, which cannot be adequately addressed using single-dimensional criteria such as grey-scale gradients. Therefore, manual segmentation is necessary to ensure high accuracy, particularly in complex or ambiguous regions with noise, low contrast, or irregular structures.[34, 35] The criteria for manual segmentation were based on the unique structural features of the identified organelles. Chloroplasts: Typically rounded or oval, containing densely packed thylakoid membranes that form stacks called grana.[36] Starch Granules: Spherical or ellipsoidal, varying in size, and present only inside chloroplasts.[37] Mitochondria: Elongated or oval-shaped, smaller than chloroplasts, moderately electron-dense, and often located near the cell periphery or other organelles.[38] Vacuole: The largest organelle, less electron-dense, centrally located, and containing cell sap.[39] Stromules: Thin, tubular structures originating from plastids.[40] Chloroplast protrusions: Similar with stromules, but the structure with short and large stromal area or tubular projections, the status just before detached from the main chloroplast body.[41]

For segmentation, selected cell organelles that were completely captured in the examined guard cell section, were manually identified according to their size, shape, color, and structure. Vacuoles were identified by their relatively large size and color, grey interior and slightly darker surrounding. Chloroplasts were identified by their round shape and the contrasting, complex thylakoid structures inside. Uniform grayish areas contrasting with the dark thylakoid structures surrounded by a white border within the chloroplasts were identified as starch granules. Mitochondria were identified by their elongated, rounded shape and darker appearance compared to vacuoles. The structures were individually labeled on selected slice images on XY-plane. After the initial labeling of an object, a morphological interpolation suitable for regularly shaped spheres was used to extend manually introduced labeling for the unsegmented image slices. The interpolated labeling was then manually corrected along the Z-axis for deviations from the original structure in the individual XY-sections. From the obtained 3D objects, organelle surface area[42] and volume were exported using Dragonfly algorithms. Data were plotted using Prism GraphPad (software version 8.3.1; 332).

## 3 Results and Discussion

The aim of our study was to evaluate the advantages and applicability of cryo-LM and cryo-FIB-SEM for analyzing higher plant cells, using guard cells in *V. faba* as a model organism. As a "proof-of-concept," we focused on the optimal dataset obtained from a single stoma under controlled conditions.

### 3.1 CLEM guided Cryo-FIB-SEM volume imaging for Guard Cell In Depth Ultrastructure Analysis

Cryo-Correlative Light and Electron Microscopy offers significant advantages over Integrated Light-SEM Systems.[43–46] Correlative light microscopy provides higher resolution and allows easy switching between magnifications by changing the objective lens, enabling both overview imaging for navigation and high-resolution fluorescence data for FIB-SEM analysis. Integrated Light-SEM systems, though convenient, can suffer from misalignment due to challenges like mechanical drift, resolution disparities, and calibration issues, making them less versatile for diverse biological samples and experimental setups. In contrast, Correlative Light-SEM systems, while more labor-intensive, provide greater flexibility and accuracy in aligning images from the two modalities. For these reasons, we utilized correlative light and FIB-SEM systems in conjunction with cryo-specimen preparation in our study.

Light/Fluorescence Microscopy (L/FM) serves the dual purpose of assessing sample quality (e.g., sample integrity and the presence or absence of a very thick (>> 1µm) ice layer on top of the epidermis surface) and facilitating navigation to areas of interest for subsequent FIB-SEM data acquisition. Cryo-LM reflection or fluorescence imaging with ZEN provides an overview of the sample, as illustrated in Figure 2 (A-C), showcasing clear intact stomata/guard cells within the leaf epidermis without visible ultra-thick ice contamination. Additionally, the autofluorescence of chloroplasts with a maximum signal around 638 nm not only confirms the vitality of guard cells but also indicates the approximate positions of chloroplasts within them (Figure 2 D-F). The chloroplast autofluorescence can be used as a rough proxy for chloroplast thylakoid membrane integrity,[47] because the signal becomes weaker as a result of thylakoid disassembly such as in the case of plants exposed to chloride stress.[48] This process helps to choose a representative stoma from the epidermal peel. Furthermore, under the fluorescent mode, the number of chloroplasts per guard cell can be visualized, as shown in Figure 2 D and F, indicating a total of 15 chloroplasts per pair of guard cells, such as eight for the left one and seven for the right one in diploid *V. faba* plants. The number of chloroplasts in guard cells is largely dependent on the ploidy level, with diploid plants typically having 11 to 15 chloroplasts in a single stoma.[49]

Cryo-LM imaging enabled the easy localization of native stomata on the epidermis attached to the grids, facilitating the observation of cell morphology (see Figure 2 A-C). Subsequently, samples were transferred to the cryo FIB-SEM for volume imaging. To avoid charging of the sample surface and to allow artifact-free SEM imaging, the sample has been sputter coated with platinum inside the Quorum prep chamber (attached to the Crossbeam chamber). The positive effect of the sputter coating layer is demonstrated in Figures 3A-D. Figures 3A and 3B show the cell surface before and after sputter coating, with charging artifacts highlighted by a dotted orange borderline in Figure 3A. Figures 3C and 3D depict two stomata: one with charging artifacts, particularly around the stoma rim (orange arrow in C), and the other without charging (D). The avoidance of charging is not only important for surface imaging, but also to precisely target the area of interest with the ion beam during FIB milling of a trench (Figure 3 E) and for the subsequent 3D FIB-SEM data acquisition, as shown exemplary in a single raw SEM image from a 3D FIB-SEM dataset (Figure 3 F).

The raw FIB-SEM images are impaired by a visible curtaining effect (Figure 4 A). They could be efficiently “destriped” by applying the VSNR algorithm implemented into the ZEN software yielding clear and contrast rich images of the guard cell (Figure 4 B). The same software allowed us to inspect the 3D dataset in the three XY, XZ and YZ image planes using the orthoview representation (Figure 4 C). The dataset for this study is contextualized by overlaying fluorescent light microscopy (LM) data with SEM images and FIB lamellae, after targeting the area of interest in the FIB-SEM (Figure 4D). Based on the processed dataset subcellular regions could be identified (Figure 5). Subsequently, the 3D reconstruction was used to build models of the various cell organelles to analyze their morphology and spatial distribution (Figure 6).

**Figure 5.**
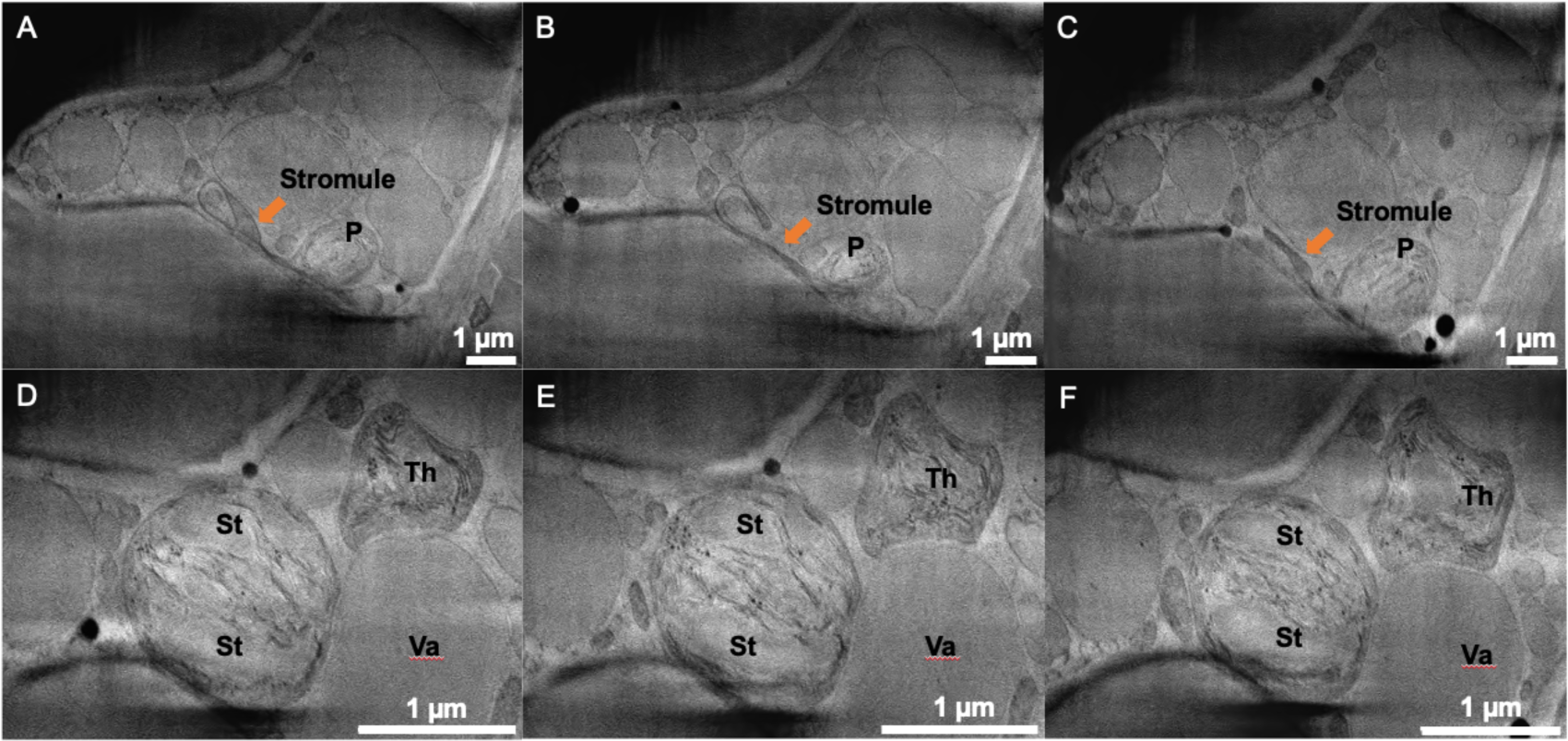
Identification of cell organelles in processed cryo-FIB-SEM images. A-C: Stromules with different morphology, in contact with either plastids or other organelles, as indicated by orange arrows; P, plastid. D-F: Identification of different organelles: Th, thylakoid; St, starch granules; Va, vacuole.

**Figure 6.**
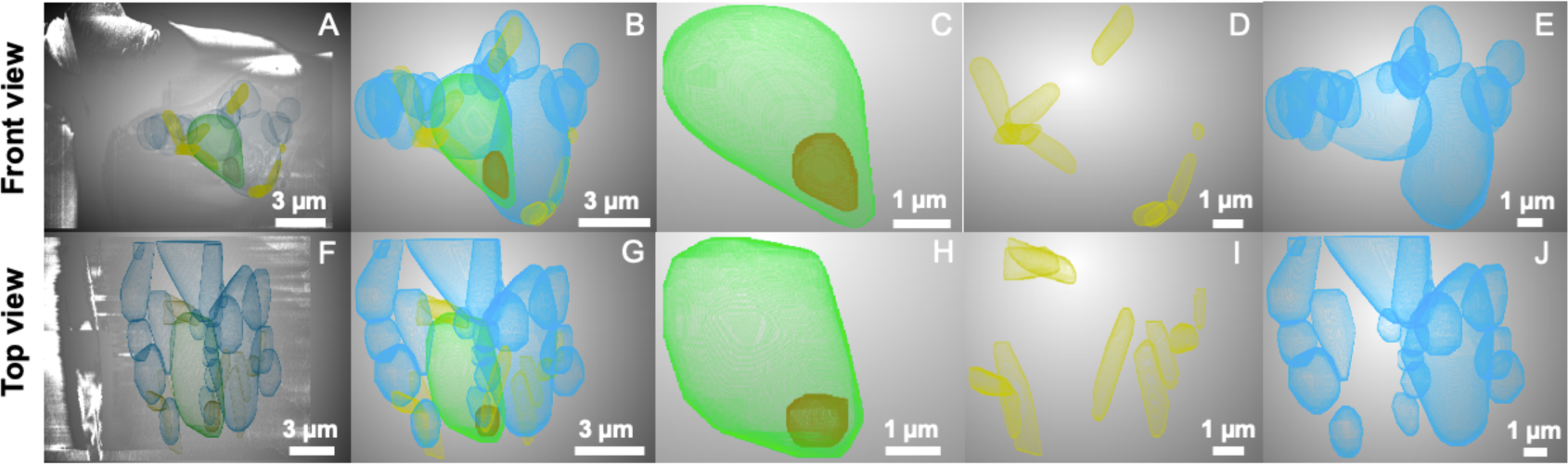
Morphology, topography, and spatial arrangement of organelles illustrated as 3D models of the reconstructed cryo-FIB-SEM dataset. Front view: A-E; A, XY plane through the 3D FIB-SEM dataset and the corresponding organelles; B, overlay of 3D morphology of chloroplasts (green), chloroplast protrusion (orange), mitochondria (yellow), vacuoles (blue); C, chloroplasts (green) and chloroplast protrusion (orange); D, mitochondria (yellow); E, vacuoles (blue); Top view: F-J; F, XZ plane through the 3D FIB-SEM dataset and the corresponding organelles; G, overlay of 3D morphology of chloroplasts (green), chloroplast protrusion (orange), mitochondria (yellow), vacuoles (blue); H, chloroplasts (green) and chloroplast protrusion (orange); I, mitochondria (yellow); J, vacuoles (blue).

As stomata play a central role in plant physiology by regulating leaf transpiration and CO2 uptake, understanding their biochemical and structural features, particularly at the subcellular level, is crucial. In a recent study involving *Arabidopsis thaliana*, 10-day-old primary leaves were fixed using paraformaldehyde/formaldehyde/osmium tetroxide in cacodylate buffer, dehydrated in a graded ethanol series, and embedded in propylene oxide and Quetol 651 resin (a low viscosity water-miscible epoxy resin) to visualize guard cell plastid morphologies and organelle ultrastructure.[50] Various organelles were observed, including the nucleus, vacuoles, chloroplasts, starch granules, mitochondria, plastoglobules, and stromules.[50] However, the chemical fixation-resin embedding process poses a significant risk of altering the native state of leaf epidermis, particularly affecting sensitive structures like stromules due to their responsiveness to environmental changes. Chemical fixation-resin embedding usually takes more time than cryo-immobilization by plunge freezing, thereby increasing the likelihood of unintended modifications. Furthermore, microscopic observation of stromules in freshly detached leaves can also lead to state alterations. In contrast, rapid cryo-immobilization enables the visualization of dynamic changes in stromule states by comparing different time points during sample preparation. For instance, stromules have been shown to rapidly extend or retract within minutes after detachment from the plant or in response to stimuli.[51] In many cases, green fluorescent protein (GFP)- modified plants facilitate easy observation of stromules under a fluorescent or confocal microscope,[52] whereas natural plants necessitate EM, posing challenges in sample preparation. High-pressure freezing and freeze substitution techniques have been employed to fix leaf cells and successfully observe the 2D structure of stromules in rice plants without genetic modification.[13] This sample preparation method successfully avoids the use of chemical fixation such as PBS solution or aldehydes.[13] However, embedding still involves the use of hazardous heavy metal compounds like osmium tetroxide, requiring a freeze substitution process ranging from a few hours to 88.5 hours.[13, 53] Moreover, high-pressure freezing has been reported to potentially disrupt the plasma membrane and cell wall and may not preserve chloroplast morphology as effectively as cryo-immobilization via plunge freezing alone.[13]

In our study, cryo-fixation achieved by plunge-freezing in liquid ethane could be completed within minutes without the need for chemical fixation and embedding. Stromules were clearly visualized in the serial volumes obtained (see Figure 5 A-C, Video S1), revealing not only their origin from the plastid but also their ability to extend and make contact with other organelles. Based on these images (Figure 5 A-C), the average diameter (0.36 μm) and length (3.33 μm) of stromules (n=3) were measured and the results matched with the literature indicating 0.35–0.85 μm in diameter and of variable length, from short beak-like projections to linear or branched structures up to 220 μm long in tomato fruit mesocarp cells and GFP-based modified plastids of tobacco and petunia plants.[54, 55] Furthermore, the volumetric FIB-SEM data enabled the generation of a 3D model illustrating the spatial relationships of chloroplast and chloroplast protrusion (CP) (see Figure 6C and H, Video S2). Stromules and CPs are tubular projections including stroma without thylakoid membranes. Generally, stromules are the structure with long and thin tubules. On the other hand, CPs are the structure with short and large stromal area or tubular projections which could be the structure just before the separation from the main chloroplast body.[41] Then we discussed the CP (n=1) volume (1 µm^3^) and surface area (5 μm^2^) with the literature indicating the volume of CP blows 2 µm^3^. [41] Also the feret diameter of CP was measured between 1 and 2 μm in parallel with the reported data of up to 5 μm.[56] Furthermore, the reconstructed 3D model provides a comprehensive visualization of the cell and organelles from two perspectives: a front view (see Figure 6A to E) and a top view (see Figure 6F to J), facilitating the analysis of the spatial arrangement of organelles such as chloroplasts, CPs, mitochondria, and vacuoles within the guard cell (see Video S1 to S5). These insights can be further extended by extracting number, surface area, and volume information from the 3D model generated from the volumetric FIB-SEM data. Consequently, these near-to-native quantification results will contribute to a deeper understanding of plant cell responses to environmental factors. For instance, the dynamic morphology and tubular structures of stromules and CPs play a crucial role in signal and macromolecule transfer between plastids and organelles, particularly under varied abiotic (e.g., osmotic stress, high light) and biotic stresses (e.g., pathogens).[41, 57]

### 3.2. Revealing Guard Cell Organelle Properties: Insights from Cryo-FIB-SEM and CLEM Analysis

Although previous studies have reported stomatal size and volume based on 3D model reconstruction,[58] organelle surface area and volume were not included. In this study, we have successfully determined the spatial arrangement, volume, and surface area of individual organelles of *V. faba* guard cells in a near-to-native state (see Figure 6 and 7).

**Figure 7.**
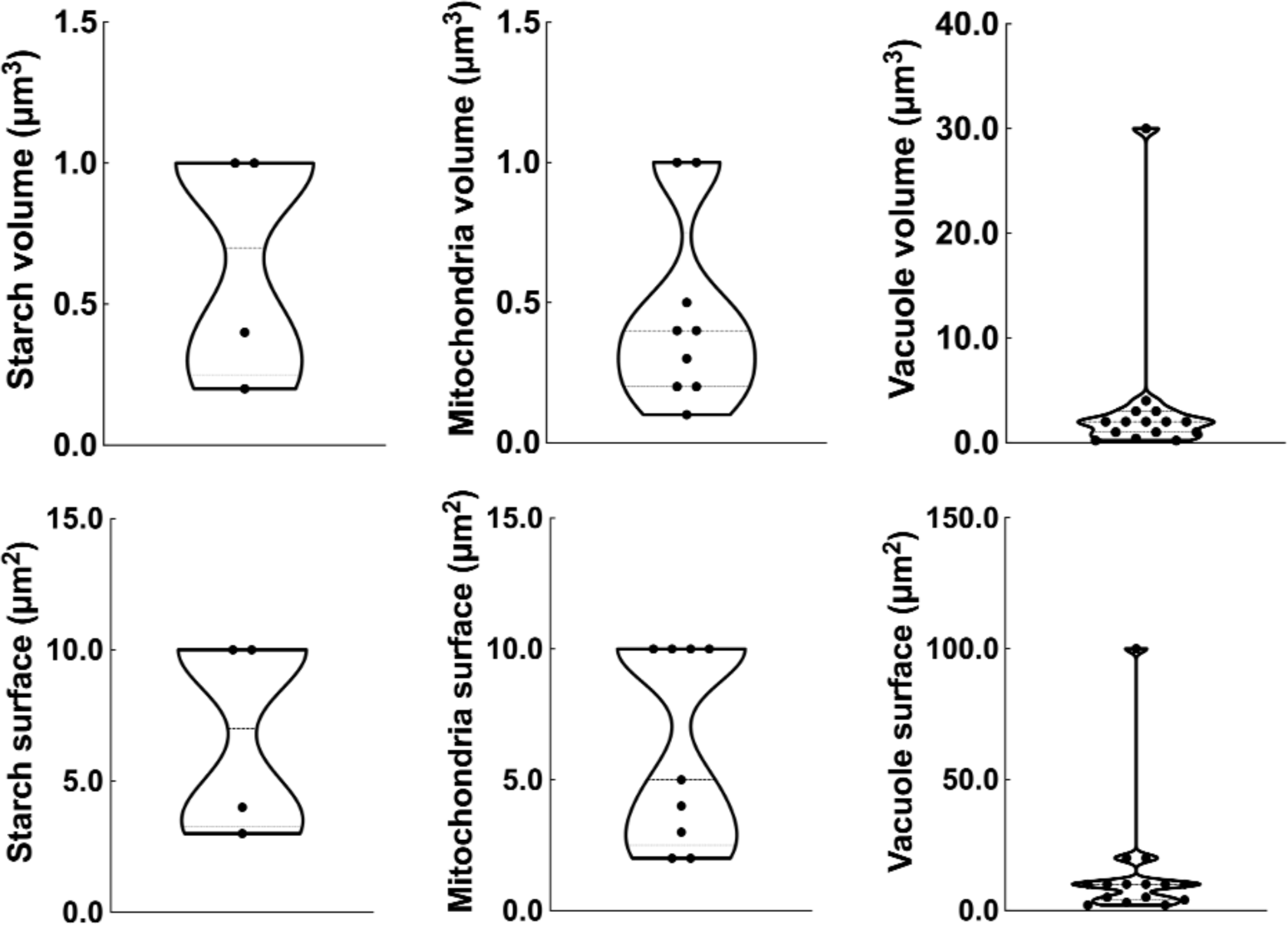
The quantified volume and surface area of the identified guard cell organelles based on reconstructed 3D models. The number of raw data replicates, indicated by black dots, depends on the number of organelles identified in the analyzed guard cell section. Starch granules in a chloroplast, n=4; Mitochondria, n=9; Vacuoles, n=15.

In this study, one guard cell was selected with the best quality to proceed the organelle quantification. The reconstructed chloroplast (n=1) (Figure 6 C and H, Video S2) had a volume of 20 µm³ and a surface area of 100 µm², fitting within the FIB-SEM- produced chloroplast volume range of 15 to 90 µm³ and surface area range of 30 to 180 µm² in rice mesophyll cells.[59] More importantly, the accuracy of quantifying chloroplast size using the FIB-SEM technique has been validated by TEM data.[59]

Additionally, our chloroplast volume (20 µm³) was much lower than previously reported data, such as 68.2 µm³ in wheat,[60] 52.5 µm³ in rice,[59] and 22.4 µm³ in chickpea.[60] This discrepancy is reasonable because guard cell chloroplasts are typically smaller than mesophyll chloroplasts.[61]

The size of starch granules plays a vital role in plant responses to external stimuli, as starch metabolism regulates stomatal aperture.[62–64] Consequently, accurately quantifying starch size in guard cells is essential. In our study, we successfully quantified starch granule size (n=4) using a reconstructed 3D model (Figure 7, Video S3). Results showed an average volume of 0.65 µm^3^, a surface area of 6.75 µm^2^, and a surface area/volume ratio of 10.38 µm^-1^, which were within the reported range of starch volume (0-20 µm^3^), surface area (0-130 µm^2^), and surface area/volume ratio (3-35 µm^-1^) in Arabidopsis leaves.[65] However, sample preparation methods can cause differences in results, even when using the same FIB-SEM technique. Chemical resin- embedded samples revealed significantly smaller starch granule volumes (0.12 µm^3^) and surface area-to-volume ratios (2.1 µm^-1^) in chloroplasts of cell-cultured algae compared to our data (0.65 µm^3^ for starch and 10.38 µm^-1^ for the ratio) from cryo-fixed guard cell chloroplasts of *V. faba*.[25] Besides, the plant itself might be another inherent reason for the difference, because algae (*Chlorella pyrenoidosa* cells) is a kind of avascular lower plant, while *V. faba* is a “real” vascular higher plant.

The average volume and surface area of mitochondria (n=9) (Figure 6 D and I, Figure 7, Video S4) were found to be 0.46 µm³ and 6.22 µm², respectively. These measurements align closely with previously published literature, which reports an approximate mitochondrial volume of 0 to 4.5 µm³ and a surface area of 0 to 33 µm², based on 250 reconstructed 3D mitochondria models in Arabidopsis mesophyll cells.[66] Our study found that the average volume and surface area of vacuoles (n=15) (Figure 6 E and J, Figure 7, Video S5) in the guard cells of *V. faba* were 3.59 µm³ and 14.73 µm², respectively. These volumes are notably lower than the range of 30 to 240 µm³ reported in the first leaf primordia of Arabidopsis.[67] This demonstrates the considerable variability in cellular structures observed not only between different plant species and developmental stages, but also in relation to the effect of stomatal aperture. Stomatal closure can result in the formation of numerous small vacuoles due to guard cell shrinkage.[68, 69]

In brief, the results of the preliminary quantification of cellular organelles using the cryo-FIB-SEM tomography workflow appear reliable, as they align with data reported in the literature. Under this assumption, the cryo-FIB-SEM system offers advantages because it preserves plant samples in a near-to-native state. However, improvements are needed in the workflow for processing and segmenting serial image data, particularly in automating organelle identification using machine learning approaches to achieve reproducible and efficient segmentation of larger datasets.[70] The quantification of authentic volumes and surface areas of individual subcellular organelles within guard cells represents a significant advancement, especially in studying plant responses to abiotic and biotic stresses. Factors such as nutrient deficiency or toxicity, drought, heat stress, high temperatures, salt stress, and pathogen diseases can induce variations in organelle structure within guard cells.[71] Understanding these variations not only illuminates the impacts of external stressors but also reveals the cellular mechanisms plants employ to adapt to challenging environments.

## 4 Conclusion

A tentative application of correlative cryo-LM and cryo-FIB-SEM volume imaging was demonstrated to be feasible for the first time on higher plant guard cells. Cryo fixation via plunge freezing is a rapid alternative to lengthy freeze-substitution protocols, and preserves samples close to their native state. Correlative cryo-LM and FIB-SEM microscopy facilitates easy navigation and detailed visualization of subcellular organelles within guard cells. FIB milling allows for the cross-sectioning of single cells, generating volumetric data sets. After reconstructing serial image data, organelle structures can be analyzed in detail. This workflow is particularly advantageous for plant physiological research on environmental influences on plant cell structures. It is especially useful for studying CPs and stromules, which are very sensitive to sample preparation and highly susceptible to structural modifications. Understanding these structures is vital for comprehending plants’ environmental responses at the cellular level. Cryo-FIB-SEM has the unique advantage of maintaining their native state through rapid plunge freezing, eliminating the need for preparatory chemical treatments. This method also facilitates potential elemental mapping. Integrating biomineral monitoring detectors, such as highly sensitive Energy Dispersive X-ray (EDX) or Energy selective Backscattered (EsB)[72], into the cryo-FIB-SEM system can map elemental distributions at the cellular and subcellular levels, further enhancing the understanding of cellular ion relations and plant cell responses to mineral applications.

Overall, our study shows that correlative cryo LM and FIB-SEM volume imaging can be used to investigate higher plant cells in their near-to-native state. This method will facilitate quantitative research to understand the reaction of plants to environmental stress such as salinity and drought on the ultrastructural level.

## Acknowledgement

The work was financially supported by the Deutsche Forschungsgemeinschaft (DFG), project number 491678431. We thank Katharina Hipp for her assistance with the plunge freezing of leaf epidermal fragments. Special thanks to Yiwei Gao, a master’s student at the Macro Agriculture Research Institute, College of Resources and Environment, Huazhong Agricultural University, for his contributions to graph creation and formatting.

## Data Availability Statement

The data that support the findings of this study are available from the corresponding author upon reasonable request.

## Supplementary video legends

Video S1 The spatial arrangements of chloroplast and stromule within a single guard cell

Chloroplast and stromule were represented by the large green ball and small orange ball, respectively.

Video S2 The spatial arrangements of chloroplast and chloroplast protrusion within a single guard cell

Chloroplast and chloroplast protrusion were represented by the large green ball and small orange ball, respectively.

Video S3 The spatial arrangements of chloroplast and starch within a single guard cell

Chloroplast was represented by the large green ball, starch granules by four small purple balls, and the orange ball designated chloroplast protrusion.

Video S4 The spatial arrangements of mitochondria and vacuoles within a single guard cell

Mitochondria were represented by orange sticks of varying lengths.

Video S5 The spatial arrangements of vacuoles within a single guard cell

Vacuoles were represented by blue balls of different sizes.

---

**Figure S1.**
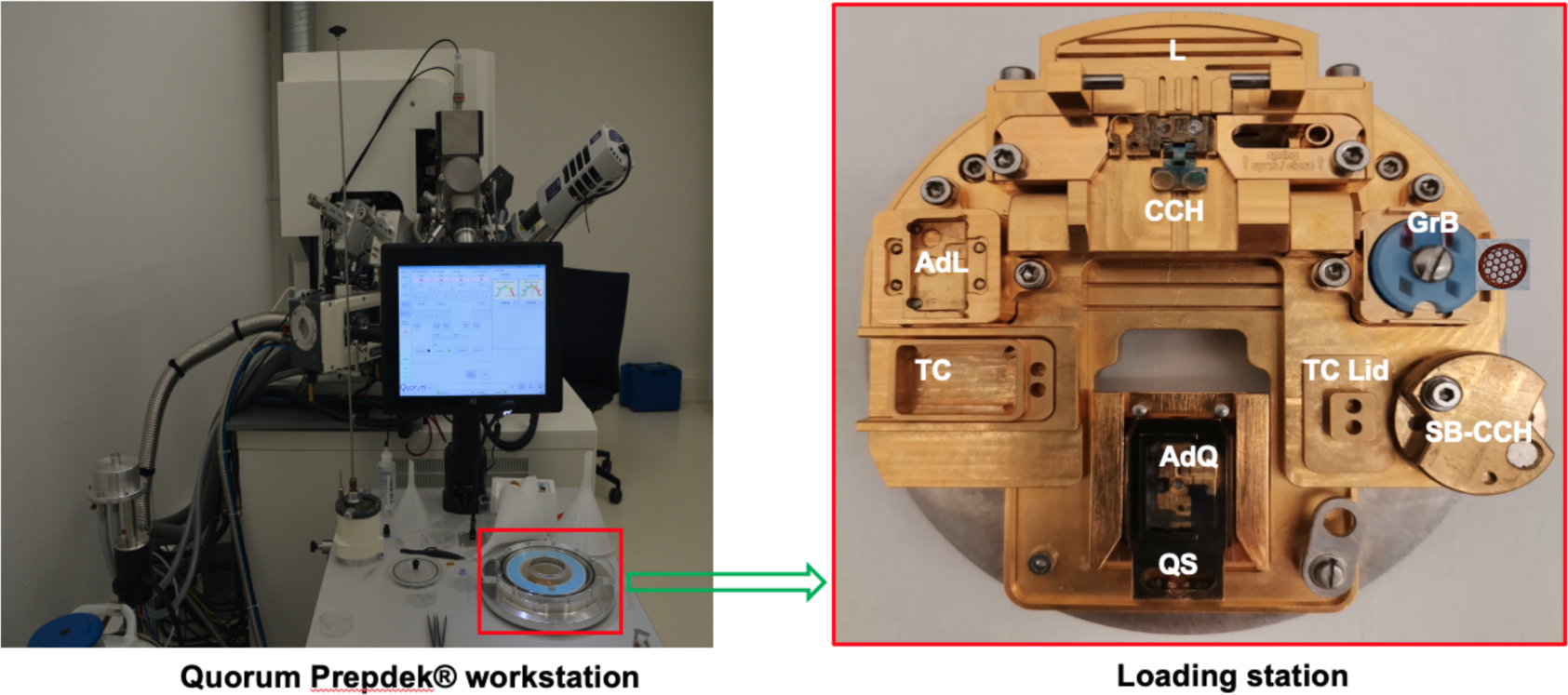
Details of loading station in the Quorum Prepdek® workstation. The loading station has dedicated positions for mounting and transferring samples under liquid nitrogen conditions. Plunge-frozen samples on TEM grids typically arrive in a grid box (GrB). Using fine tweezers, the TEM grid is transferred to the loading position (L) from where it can be gently pushed into the correlative cryo holder (CCH) - two grids can be mounted at a time. For transfer into the LM the CCH is mounted onto the Linkam cryo-adaptor (AdL), and temporarily stored in the transfer container (TC) with the TC lid sealing the container. After LM imaging the sample inside the CCH is moved back to the loading station where it is transferred onto the Quorum shuttle (QS). Using the Quorum transfer device, the sample can now be transferred into the Zeiss Crossbeam 550 for cryo-FIB-SEM data acquisition. After the experiment, the sample(s) in the CCH can be stored in a storage box (SB-CCH).

